# EcoToxFred: A knowledge-graph-grounded conversational agent for exploring ecotoxicological monitoring and hazard data

**DOI:** 10.1101/2025.07.04.663152

**Authors:** Patrick Scheibe, Thilo Simon Schmid, Jana Schor

## Abstract

Assessing chemical risks to aquatic ecosystems requires the joint interpretation of monitoring data, ecotoxicity data, and derived risk indicators, yet such integrated data are rarely accessible to users without database expertise. We present EcoToxFred, an open-source conversational agent that provides natural language access to a curated knowledge graph of chemical monitoring and hazard data for European surface waters. EcoToxFred couples a large language model (LLM) with a Neo4j graph database in a retrieval-augmented generation (RAG) architecture. An agent interprets user questions, selects tools, translates questions into graph queries, and returns answers as text, tables, or interactive maps, together with the executed query for reproducibility. We evaluated EcoToxFred with GPT-4.1 and GPT-5 on a benchmark of 25 questions, including six questions derived from an expert workshop, each posed three times. Answers were scored for tool selection, execution accuracy against human-written reference queries, and goal accuracy using the RAGAS evaluation framework with an independent LLM judge. With both models, EcoToxFred selected an appropriate tool in at least 92% of runs and reproduced the reference result in 74% of runs with a reference query, including all lookup questions. Goal accuracy was 52% with GPT-4.1 and 63% with GPT-5. Failures concentrated on multi-step analytical questions, incomplete presentation of long results, and mismatches between user and database vocabulary. EcoToxFred shows how knowledge-graph-grounded LLM agents can lower the barrier to ecotoxicological data exploration while keeping answers traceable to the underlying data, and its architecture is transferable to other ecological data infrastructures.

## 1 Introduction

Pollution, notably chemical pollution, is one key element of the triple planetary crisis, alongside climate change and biodiversity loss (Sigmund et al. 2023). These crises are profoundly interconnected and demand integrated data approaches supporting cross-cutting environmental assessments and decision-making processes.

Reliable ecotoxicological data analyses are essential in environmental decision-making because they are crucial for assessing the impact of chemical pollutants, e.g., on aquatic ecosystems, and supporting regulatory risk assessments of chemicals. Environmental monitoring data on chemical concentrations must be integrated with laboratory-based toxicity data for aquatic organisms to evaluate the ecological effects of chemical substances. Such tests capture biological responses at different levels of organization and provide the foundation for linking chemical exposure to adverse outcomes (Schuijt et al. 2021). Species sensitivity distributions (SSDs) enable prioritization and risk management of chemicals in line with regulatory mandates such as the European Water Framework Directive (Posthuma et al. 2019). Biomarkers and bioassays are increasingly used to detect sublethal pollutant effects under real-world conditions (Connon et al. 2012), and standardized methods support cross-study comparability and regulatory application (Rosner et al. 2024). Furthermore, the structured and reproducible evaluation of ecotoxicity data plays a key role in deriving safe concentration thresholds and ensuring robustness in regulatory contexts (Moermond et al. 2016).

Accessing and analyzing eco-toxicological data remains challenging despite the growing availability of environmental data, especially for users without expertise in database technologies. Traditional relational databases often lack the flexibility needed to capture the complexity of environmental systems, including non-linear and nested relationships among chemicals, species, and habitats (Yang and Neagu 2013). These systems generally require knowledge of structured query languages (e.g., SQL), limiting usability for non-computer scientists (Labrinidis and Jagadish 2012). Moreover, environmental datasets are frequently heterogeneous and dynamic, making them poorly suited to static tabular storage models (Wang and Xu 2018). As a result, non-computer scientists and key stakeholders, including regulators and policymakers, face barriers in engaging with and interpreting the data required for evidence-based decision-making (Ginsberg et al. 2019; Attaf et al. 2018).

Graph databases offer a flexible solution for implementing ecotoxicological knowledge graphs. They provide a powerful means to model the complex interconnections typical in ecotoxicological data by representing entities such as chemicals, species, effects, and environmental compartments as nodes, and their interactions as relationships, supporting intuitive and semantically rich data structures (Olken 2003; Memarzadeh et al. 2018). These databases are well-suited to integrate diverse datasets, accommodate changes in schema, and support exploratory querying at scale (Dominguez-Sal et al. 2010; Junghanns et al. 2017). More recent approaches that demon-strate the usage of graph databases in scientific data integration are, for example, the `KnowWhereGraph` (Zhu et al. 2025), a large–scale geo-knowledge graph for interdisciplinary knowledge discovery and geo–enrichment. `SustainGraph` (Fotopoulou et al. 2022) was developed to track progress towards the United Nations Sustainable Development Goals at national and regional levels. Graph databases’ dynamic representations and relationship-centric data storage offer a promising foundation for ecotoxicological decision-support systems.

Recent advances in large language models (LLMs) and retrieval-augmented generation (RAG) (Lewis et al. 2020) have expanded human-AI interaction by unlocking new possibilities for accessing and interacting with complex data sources. While LLMs excel at interpreting and generating human-like language, they do not possess true understanding and cannot access up-to-date or domain-specific knowledge. RAG addresses this limitation by dynamically grounding model outputs in external, structured sources such as databases or knowledge graphs. This hybrid approach improves accuracy, contextual relevance, and transparency. LLM systems that are augmented with knowledge graphs were shown to drastically reduce the query efforts and hallucinations (see, e.g., Silveira et al. (2024)). Especially in scientific domains, grounding LLMs with curated knowledge resources offers an opportunity to support expert-level dialogue while maintaining accessibility for non-specialists (Stahl 2023; Brown et al. 2020). RAG-based systems have demonstrated value in other domains. Tools like `MedRAG` (Zhao et al. 2025), `ChemCrow` (Bran et al. 2024), `HazardChat` (Silveira et al. 2024), `ChatClimate` (Vaghefi et al. 2023), and `LegalBench` (Pipitone and Alami 2024) are transforming access to knowledge in medicine, chemistry, climate science, and law. While the emerging `HazardChat` platform addresses chemicals and their hazardous effects, it focuses on linked hazard data. Applications in environmental sciences that involve exposure, hazard, and, therefore, risk analyzes remain underexplored.

We introduce EcoToxFred, a domain-specific, retrieval-augmented chat agent that bridges LLMs with a graph database containing integrated chemical and ecotoxicity data from European surface waters. It enables users to query and explore environmental knowledge through natural language, removing the barrier of technical query syntax. The system uses a decision-making agent to orchestrate tool usage, transforming questions into structured queries and refining outputs before delivering them back to the user. EcoToxFred supports output formats ranging from text to tables and interactive geographic maps, making the exploration of environmental knowledge intuitive and flexible. While designed for ecotoxicology, the approach generalizes to other scientific domains reliant on structured data and expert-level interpretation. The impact potential of EcoToxFred is high: EcoToxFred would significantly lower barriers for policy-makers, researchers, and non-technical stakeholders to access and interpret environmental data.

Integrating large language models with graph-based knowledge systems represents a promising direction for responsible, human-centered AI interaction, improving science communication and decision support.

## 2 Results

We have developed EcoToxFred, a react-agent to interact with domain-specific complex data via a user-friendly browser interface. The agent leverages a large language model (LLM) in a tool-oriented architecture. The LLM is grounded with data from a knowledge graph stored in a graph database. Retrieval-augmented generation (RAG) is used to retrieve and incorporate information into the discussion between EcoToxFred and the user. This approach provides precise, domain-specific, context-aware responses based on common and expert knowledge successfully.

The available tools include `CypherSearch` and `GeographicMap`, which transform the user query into a Cypher graph querying language statement, submit it to the graph database, evaluate the results of these database requests, and transform them back into a human language response to the user. `CypherSearch` is triggered by general requests to the data, such as user queries that contain triggers like ‘How many,’ ‘List,’ ‘ Show relationships,’ or ‘Which species.’ `GeographicMap` is activated whenever the user requests to show data on a map. Example triggers in the user query are: ‘Where,’ ‘Show,’ ‘Show map,’ ‘Visualize sites,’ or ‘Map of,’ or geographical regions like river or country names. The third tool `WikipediaSearch` allows responding to more general questions that do not directly require accessing the data holding the ecotoxicological information, such as information about a specific chemical, place, or metric. Examples of triggers for this tool are ‘What is,’ ‘Explain,’ ‘Overview,’ or ‘General info.’ General conversation is handled by the LLM directly, without a tool call.

The agent’s behavior and the orchestration of its tools build the core of EcoToxFred, governed by a ReAct (reasoning and acting) architecture (Yao et al. 2022). This agent dynamically reasons about user queries, selects appropriate tools, and iteratively refines responses in a memory-aware workflow, see Figure 1. Each interaction begins with the LLM decomposing the user input and deciding on the required tools and reasoning steps, e.g., a compound-related query may require using both `WikipediaSearch` for context and `CypherSearch` for factual, structured data. Tool outputs are passed back into the agent loop, and independent tool calls can be executed in parallel. The workflow supports conditional branching and a retry logic. If a tool fails to return a valid result, e.g., due to query syntax issues or empty result sets, the agent re-evaluates the internal state and can retry queries with modified parameters or report. This procedure ensures robustness against ambiguity and improves response coverage for under-specified queries.

**Fig. 1:**
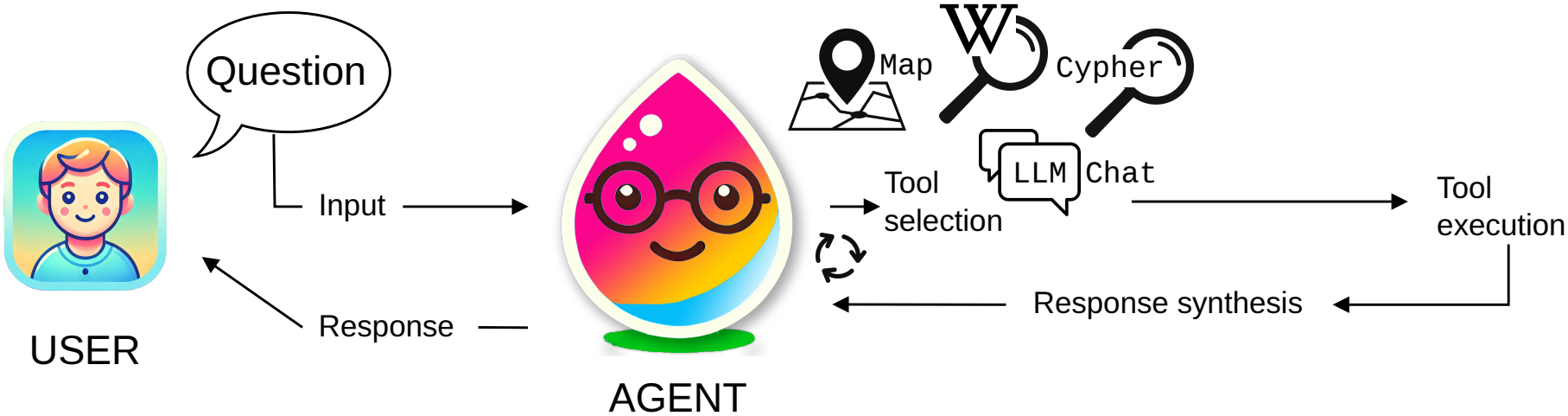
The agent’s workflow follows a reasoning-and-acting loop: it selects tools, observes their results, and either calls further tools or synthesizes the response. “LLM Chat” denotes the case in which the system answers directly without calling a tool.

We quantify how reliably this orchestration mediates between the LLM and the structured data in the bench-mark evaluation below (Section 2.1).

Use cases demonstrate the capabilities of EcoToxFred through three representative user queries (see Figure 2) that reflect expected questions from researchers or stakeholders in the context of ecotoxicological data exploration. Each use case illustrates how EcoToxFred interprets a natural language query, selects appropriate tools, executes the necessary data retrieval steps, and formats the response. See Figures 2a–2c for the use case results described below.

**Fig. 2:**
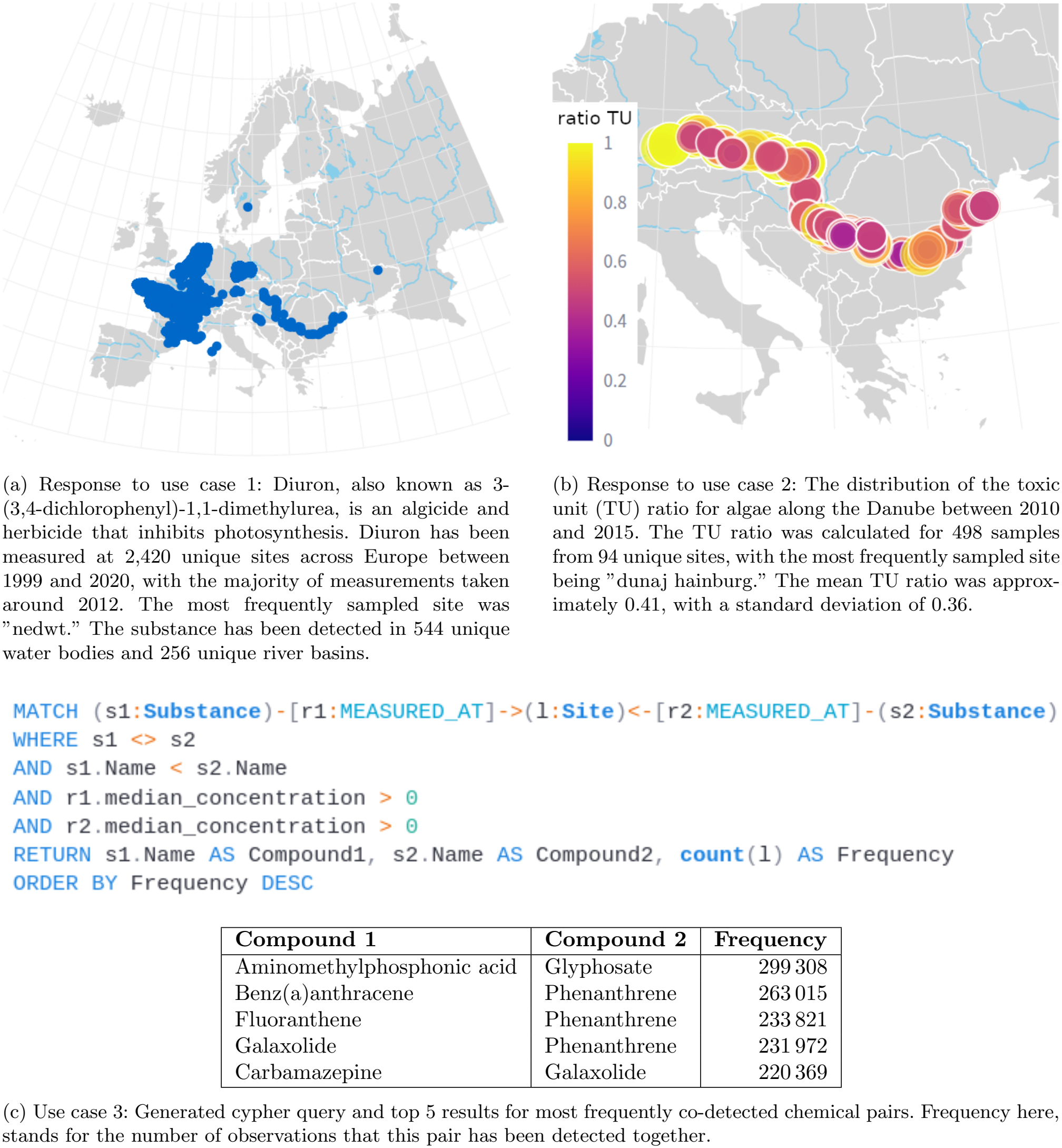
Overview of EcoToxFred’s system components and capabilities. The captions for subfigures (a) and (b) are generated by EcoToxFred itself (shortened) as part of the response to user queries.

### Use Case 1: Querying Chemical Occurrence and Background Information

> *“What is Diuron and where has it been measured?”*

The agent decomposes the question and routes it to two tools: (i) `WikipediaSearch` to retrieve general information on Diuron as a compound, and (ii) `GeographicMap` to query the graph database for locations where Diuron has been measured in European surface waters. The result (see Figure 2a) includes an interactive map of Europe showing all sites where Diuron has been measured at least once. The user may zoom in and hover over the data to retrieve more information about the individual measurements. Further, the result includes a summary of Diuron’s use and mode of action from Wikipedia, as well as details about the number of sites, rivers/lakes, river basins, the measurement time range, and the most frequently measured sampling site calculated from the retrieved data. This use case demonstrates EcoToxFred’s ability to provide both structured data and unstructured context, linking chemical identity with its environmental monitoring footprint.

### Use Case 2: Spatial Visualization of Risk Indicators

> *“Show the ratioTU distribution for algae along the Danube from 2010 to 2015.”*

The agent interprets the query as requiring a geospatial aggregation of Toxic Units (TU) for algae, filtered by time and river region. The `CypherSearch` tool is used to extract the relevant data, and the `GeographicMap` tool provides the result as an interactive geographic visualization. The interactive map shows color-coded risk levels at monitoring stations along the Danube and an appropriate caption describing the indicator, time period, and species group. This use case highlights the system’s support for multi-modal outputs and its ability to analyze complex ecological indicators in an interpretable format.

### Use Case 3: Co-occurrence Query of Frequently Detected Compounds

> *“List all compounds detected together most frequently.”*

This prompt triggers the `CypherSearch` tool, as the question requires structured access to co-occurrence patterns within the graph. The agent recognizes that the user asks for compound pairs frequently appearing in the same sampling events. To fulfill this, the LLM constructs a complex Cypher query involving a self-join on the `Substance` node, filtered to avoid duplicated combinations and aggregated by co-occurrence count. The agent generates the respective complex Cypher (Figure 2c, top) and returns the top 5 pairs as a table (Figure 2c, bottom). The response also includes a link to the associated Zenodo repository (Schor et al. 2025a) where a downloadable Docker container with the Neo4j instance is available. Technically experienced users can reproduce and extend the query on the full dataset when running it in the Docker container.

This example highlights EcoToxFred’s ability to handle multi-hop graph queries that involve pattern-based joins and aggregation. It also illustrates the system’s awareness of computational constraints and its support for exploratory scientific analysis through direct reproducibility. The use cases were recorded with an earlier release of EcoToxFred; all quantitative results below refer to the current release.

#### 2.1 Benchmark evaluation

To assess how reliably EcoToxFred answers domain-specific questions that require the knowledge it has been grounded on, we evaluated it on a benchmark of 25 questions (Section 3.6, Supplementary Table S1, questions D00–D18 and W01–W06) with GPT-4.1 and GPT-5 as agent models. Each question was posed three times in independent sessions, resulting in 75 runs per model. Each run was scored with three metrics (Section 3.6): tool selection, i.e., whether the agent called an appropriate tool; execution accuracy, i.e., whether the database queries generated by the agent returned the result of a human-written reference query; and goal accuracy, i.e., whether the final answer fulfilled the user’s goal, as decided by an independent LLM judge with the RAGAS framework (Es et al. 2024). To compare the two agent models on the same scale, goal accuracy was judged by a third model, OpenAI’s reasoning model o3 (OpenAI 2025d). Table 1 summarizes the results.

**Table 1:** Benchmark results per question category for GPT-4.1 and GPT-5 as agent models. Tool selection (T), execution accuracy (EX), and goal accuracy (G) are given in percent of runs; each question was posed three times. EX was assessed for 18 questions with a reference query or without matching data (54 runs per model; 15, 24, and 15 runs for lookup, analytical, and workshop-derived questions). G was judged by OpenAI’s o3 model for both agent models. Latency (L) is the median time in seconds from question to final answer.

|  |  | GPT-4.1 |  |  |  | GPT-5 |  |  |  |
|---|---|---|---|---|---|---|---|---|---|
| Category | Questions | T | EX | G | L | T | EX | G | L |
| General knowledge | 1 | 100 | – | 100 | 20 | 100 | – | 100 | 38 |
| Lookup and aggregation | 5 | 100 | 100 | 67 | 11 | 100 | 100 | 73 | 25 |
| Analytical | 13 | 92 | 54 | 41 | 16 | 85 | 54 | 62 | 66 |
| Workshop-derived | 6 | 100 | 80 | 56 | 14 | 100 | 80 | 50 | 82 |
| Overall | 25 | 96 | 74 | 52 | 16 | 92 | 74 | 63 | 52 |

Both models selected an appropriate tool in almost all runs and generated queries that reproduced the reference result in 74% of the runs with a reference query (40 of 54 runs each). All lookup and simple aggregation questions were answered with correct queries in every run. Analytical questions were the main source of errors, with an execution accuracy of 54% for both models, although the models failed on different questions. Typical errors were a deviating interpretation of the question, such as ranking substances by the number of affected sites instead of the number of exceedances; ignoring a dedicated relationship of the graph schema, such as the precomputed co-occurrence of risk drivers; and slightly different statistical definitions, such as alternative percentile estimators, which the strict comparison counted as incorrect. In three GPT-4.1 runs and one GPT-5 run, a generated query blocked the database and was terminated, which ended the run with an error. GPT-5 asked a clarifying question instead of querying the database when the question was ambiguous (D01), which is desirable in practice but was scored as a failure in our single-turn benchmark.

Goal accuracy was lower than execution accuracy for both models (52% for GPT-4.1 and 63% for GPT-5). A frequent reason was that the agent retrieved the correct data but presented only part of it: the `CypherSearch` tool passes only the first five rows of a result to the LLM, so answers to questions that ask for ten items or for a complete list of sites remained incomplete. The difference between the two models was largest for analytical questions (41% versus 62%), whereas GPT-5 took about three times longer to answer (median 52 s versus 16 s). Goal accuracy depended on the judge: when each model was judged by the other model instead of o3, goal accuracy was 48% for GPT-4.1 and 68% for GPT-5. When the same judge assessed the same runs in a second, independent judging pass, its verdicts agreed with the first pass in 89% (GPT-5 as judge) and 92% (GPT-4.1 as judge) of the runs.

#### 2.2 Expert feedback

We presented EcoToxFred in the workshop “CITE Meets AI: Working with LLMs, EcoToxFred, and WISE” for members of the UFZ research unit Chemicals in the Environment (CITE). Participants explored the application freely and submitted anonymous free-text feedback through an online poll (15 responses from 8 participants). Participants posed demanding scientific questions, such as identifying chemicals that are at least ten times more toxic to fish than to algae and crustaceans, which succeeded after the question had been rephrased. The feedback also revealed recurring failure modes of the release used at that time: (1) questions phrased with abbreviations or chemical classes instead of full substance names were not resolved; (2) filters on categorical properties, such as use classes, succeeded only when phrased in the exact vocabulary of the database; (3) questions on modes of action (e.g., estrogenic compounds) could not be answered from the knowledge graph, which does not annotate modes of action, and the agent incorrectly concluded that no such compounds had been measured; and (4) a request for the underlying data sources returned an incomplete list. In one case, a reported failure turned out to be a correct answer: the knowledge graph does not contain sites in France at which Iomeprol was a risk driver in 2010, whereas such sites exist in Germany. Participants further requested data upload and export, access to previous conversations, and more detailed logging. We converted the reported failures into six benchmark questions (W01–W06 in Supplementary Table S1). With the current release, the question on chemicals that are more than ten times more toxic to fish than to algae and daphnia (W04), asked in the original wording that had failed in the workshop, and the question using the abbreviation DDT (W06) were answered with correct queries in all runs of both models. Both models correctly reported that Iomeprol was not a risk driver at any French site in 2010 (W02) and retrieved the correct German sites (W01), although the answers listed only the first five of 31 sites. The question on pharmaceutical risk drivers (W03) was answered correctly in only one of six runs. GPT-4.1 searched for the use class “pharmaceutical” in lower case, whereas the database stores “Pharmaceutical”, and therefore found no drivers. GPT-5 matched the use class case-insensitively but in two runs counted all `IS DRIVER` relationships, including those that record that a substance was not a driver (`is driver = false`), which inflated the counts. For estrogenic compounds (W05), GPT-4.1 repeated the error reported in the workshop and stated that no estrogenic compounds had been measured, whereas GPT-5 searched for substance names of known estrogens and listed them, without stating that modes of action are not annotated in the knowledge graph.

## 3 Methods

### 3.1 System overview

EcoToxFred is a domain-specific conversational agent that provides natural language access to ecotoxicological data stored in a structured knowledge graph. The system is hosted as a webapp at the UFZ’s architecture and publicly accessible for user interaction via webbrowser. The system combines three main elements: a Neo4J graph database (Neo4j 2025) that stores curated and integrated environmental and ecotoxicity data, a large language model (LLM) for interpreting user input and generating responses, and a graph-based agent controller for orchestrating tool use, data retrieval, and response refinement (Lewis et al. 2020). The LLM is configurable and can be accessed through any OpenAI-compatible interface. In the public web application, users can switch between two LLM back ends: open-weight models hosted by the Helmholtz Blablador service (https://helmholtz-blablador.fz-juelich.de), which provides non-proprietary access to LLM functionality on Helmholtz infrastructure, and the proprietary GPT-5 model (OpenAI 2025a). The use cases were generated with GPT-4 (OpenAI 2025c). The benchmark evaluation was performed with GPT-4.1 (OpenAI 2025b) at temperature zero and with GPT-5 with its default sampling settings, as GPT-5 does not support a temperature parameter.

EcoToxFred (release 2026.1.0) is implemented in Python 3.13 as a ReAct agent, a multi-tool decision-making architecture (Yao et al. 2022). The agent is built with the agent abstraction of LangChain (version 1.2.6) (Chase 2022), which executes as a looped computation graph in the LangGraph runtime (version 1.0.6) (LangChain Inc. 2024). Each node in this graph represents a functional component (e.g., model call, tool execution), and edges encode transitions based on the internal state of the conversation. Access to the Neo4j graph database is enabled via the `langchain-neo4j` integration (version 0.7.0) and the `neo4j` Python driver.

The system architecture supports different output modalities. Depending on the tool and query type, responses may include plain text answers, tabular data, or interactive geographic visualizations rendered directly in the browser. Internally, all tool outputs are passed through an LLM-based response validator and formatting procedure to ensure consistency, clarity, and interpretability for end users.

### 3.2 Knowledge Graph and Data Sources

EcoToxFred builds upon a spatially curated dataset of measured chemical concentrations in European surface waters (İlhan Mutlu et al. 2025) and ecological hazard information collected and processed in (Kramer et al. 2024). In a separate project, a knowledge graph was developed that integrates these data in a knowledge graph and provides it in a Neo4j graph database (Schor et al. 2025b). The Neo4j container is publicly available at Zenodo (Schor et al. 2025a).

The knowledge graph *G* = (*V, E*) was implemented in Neo4j (version 5), a property graph database optimized for representing and traversing interconnected entities. In the graph, the nodes *V* represent key ecotoxicological and environmental concepts, specifically (1) chemicals, (2) sampling sites, and (3) species. The edges *E* capture the semantic relationships between these concepts. These relationships include the fact that a chemical was (1) measured at a particular sampling site, (2) tested for toxicity on a species, or (3) identified as a risk driver. Further, (4) there is a relation that aggregates the risk of all measured chemicals at a certain sampling site and time point for a species. Additional details, such as specific measurement values and hazard information, are stored as properties associated with both nodes and edges.

The graph is populated with

- chemical monitoring data from European surface waters; including spatial coordinates, temporal ranges, and concentration values
- ecotoxicity data such as concentration thresholds that evaluate an effect for algae, crustaceans, and fish, enabling calculation of risk indicators
- derived metrics like Toxic Units (TU) and summarized TUs per sampling site, aggregated across taxa and time periods (quarterly for the years 1998–2020) for selected regions

The graph schema supports multiscale analysis, from fine-grained sampling events to river-level summaries, and is designed for extensibility toward additional data domains (e.g., bioassay endpoints, run-off data, or land-use layers, river network data, and biodiversity). We extend this foundation in ongoing work by linking it with the EcoToxFred chat agent system, enabling interactive, conversational access to this rich dataset. All nodes and relationships are annotated with metadata such as provenance, units, time points, and coverage to support transparent interpretation and reproducibility.

The Neo4j database is securely hosted at the Helmholtz Centre for Environmental Research (UFZ) and made accessible through an internal API for Cypher-based programmatic queries and interaction with the chat agent interface.

### 3.3 The Agent

EcoToxFred employs a multi-tool, decision-making ReAct-agent to interpret user queries, select appropriate tools, and generate coherent, context-aware responses. The agent is implemented using LangChain’s agent abstraction on top of the LangGraph framework, which models computational logic as a directed graph of functional nodes. This design enables dynamic transitions between processing stages and supports persistent memory across conversation turns. A planning component maintains an explicit to-do list for multi-step requests, which the agent creates, updates, and works through while answering.

The architecture is composed of several distinct stages. Upon receiving a natural language query, the agent performs semantic analysis to infer the user’s intent. It then determines whether the request pertains to structured data (e.g., chemical concentrations, risk metrics, or site-specific measurements) and evaluates the suitability of different tools for fulfilling the query. Depending on this analysis, the agent invokes one or more tools to retrieve relevant data or generate explanatory content. These tools include interfaces for querying the Neo4j knowledge graph, accessing external textual sources, and rendering interactive maps.

LangGraph facilitates this orchestration by modeling each functional unit as a node within a computation graph. Transitions between nodes are conditioned on the evolving internal state of the agent. Memory is preserved across turns, allowing the agent to refine follow-up queries or resolve references to earlier results.

The decision-making process is guided by a prompt template library specifically designed to capture ecotoxicological reasoning. Prompt templates direct the agent to assess user needs, select the most appropriate tools, correctly format input parameters, and synthesize final responses by integrating structured outputs with contextual explanations. The overall architecture enables flexible, context-sensitive responses grounded in both domain knowledge and interactive dialogue.

### 3.4 Tool Descriptions

The EcoToxFred system incorporates a suite of modular tools, each tailored to address distinct aspects of ecotoxicological information retrieval and presentation. These tools are integrated into the agent workflow and invoked dynamically based on the user’s query characteristics.

#### CypherSearch

The `CypherSearch` tool was developed for EcoToxFred and enables structured querying of the Neo4j-based ecotoxicological knowledge graph. It builds on a modified version of the `GraphCypherQAChain` of the LangChain framework (Chase 2022) and translates natural language input into Cypher queries using a domain-specific prompt template that includes the graph schema, detailed rules, example queries, and semantic mappings for chemicals, taxa, and spatiotemporal attributes.

The tool supports both entity-centric and metric-based queries. Up to 1000 result rows are retrieved; the first five rows and summary statistics of the full result are passed to the LLM, which keeps responses within the context window. Each response is supplemented with the Cypher query used to generate it, promoting transparency and reproducibility. Queries are limited to 120 s; if a query fails or times out, the agent receives the error message and can reformulate the query. Robust input validation mechanisms handle common failure cases, such as empty or malformed queries, by providing diagnostic messages and suggestions for refinement.

#### WikipediaSearch

The WikipediaSearch tool is designed to supplement the system’s structured data with a general scientific back-ground. It extracts summaries from Wikipedia articles relevant to chemicals, ecological terms, or toxicological concepts, particularly when the graph database lacks sufficient contextual coverage. The tool uses keyword extraction and LangChain’s `WikipediaAPIWrapper` (Chase 2022) to retrieve, filter, and format relevant excerpts. The output is returned in a readable, structured format that integrates seamlessly with the system’s broader dialogue.

#### GeographicMap

The GeographicMap tool generates interactive geospatial visualizations of ecotoxicological data. It accesses site-level measurements from the knowledge graph via Cypher queries generated through LangChain’s `GraphCypherQAChain` (Chase 2022). The tool processes query results into pandas DataFrames before converting them to Plotly-compatible visualization formats. The front end renders the data using Plotly Express, supporting a range of toxicity indicators (e.g., TU, sumTU, maxTU, ratioTU) with color scales tailored to each metric.

The visualization pipeline includes preprocessing steps to validate essential geospatial fields (latitude, longitude, site identifiers) and to aggregate measurements over time or taxonomic groups. The tool enhances interpretability through hover elements displaying site names and relevant metadata such as water body, river basin, country, and temporal information. Users can explore spatial trends and hotspot regions through an interactive interface that provides both visual representation and statistical summaries of the displayed data.

### 3.5 Browser Interface and User Interaction

EcoToxFred’s user interface is implemented as a web-based application, providing a conversational experience that facilitates intuitive access to complex environmental data. Developed using the open-source Python frame-work Streamlit (Streamlit Inc. 2019), the interface supports multimodal output, allowing users to engage with the system through text queries, tabular data, and interactive geographic maps.

The main interface is composed of three components: a central chat panel for user-agent interaction, a sidebar containing example queries and system descriptions, and a response area that dynamically renders different output formats. Users may enter free-form text or select predefined queries to initiate interaction.

The system supports a range of output modalities, including:

- Natural language responses that summarize or contextualize findings.
- Tabular data that is rendered in a pandas-style format, allowing for sorting and inspection.
- Interactive maps, enabling zoom, pan, and hover functionalities for geospatial exploration.

During query processing, the interface streams intermediate updates that expose the agent’s internal reasoning. These include indications of tool selection, intermediate outputs, and reformulated subqueries. This transparency fosters user trust and supports iterative exploration of the dataset.

The interaction flow emphasizes clarity and responsiveness. Upon receiving a query, the system displays real-time feedback, including tool invocations and progress indicators. Once processing is complete, the response is rendered with context-appropriate formatting, and users are invited to refine their questions or explore related aspects via follow-up queries.

This design ensures that both domain experts and non-specialist users can efficiently navigate through and interact with ecotoxicological information, stimulating informed decision-making and interdisciplinary collaboration.

### 3.6 Benchmark design and evaluation metrics

The benchmark comprises 25 questions (Supplementary Table S1). Nineteen questions were written by the developers and cover general background knowledge (1), lookups and simple aggregations (5), and analytical questions that require multi-hop graph traversal, filtering on toxic unit thresholds, or statistical aggregation (13). Six questions were derived from failures reported in the expert workshop (Section 2.2), using the original wording of the participants where possible. One of them has no matching data in the knowledge graph and tests whether the agent correctly reports an empty result instead of fabricating an answer. Each question has a desired outcome, and 19 questions have a human-written reference Cypher query together with the answer columns that the question asks for. Where a question does not specify the number of results, the reference was restricted to the top five rows, and rows tied at a cut-off were excluded. Executing the reference queries against the knowledge graph revealed errors in four of them. A subsequent pilot run, which is not reported, revealed further mismatches between reference queries or desired outcomes and the wording of the questions, as well as an overly strict initial scoring procedure. All of these were corrected before the reported run.

The benchmark was performed separately for two agent models, GPT-4.1 and GPT-5. For each agent model, each question was posed three times, each time in a new session, to account for the non-determinism of LLM inference; this resulted in 75 runs per agent model and 150 runs in total. The query time limit described above was introduced after the benchmark; during the benchmark, queries that blocked the database were terminated externally, which ended the respective runs with an error. For every run, we recorded the full conversation including all tool calls and generated Cypher queries, the final answer, and the latency, and computed three metrics:

- Tool selection: the agent invoked at least one of the tools appropriate for the question.
- Execution accuracy, assessed for 18 questions: all Cypher queries generated by the agent in the conversation were executed against the knowledge graph, and their combined results were compared to the result of the reference query. A run was counted as correct if, for every row of the reference result, the values of the answer columns occurred together in one row of the agent’s results, after normalizing strings to lower case and numbers to four significant digits. Column names, column order, and additional columns or rows were ignored. For the question that asks for a sample of qualifying chemicals, a run was counted as correct if all returned chemicals belonged to the reference set; for the question on a typical value, either the mean or the median was accepted; and for the question without matching data, a run was counted as correct if the last generated query, on which the answer is based, returned no rows. The remaining seven questions are open-ended or combine several sub-questions and were evaluated by goal accuracy only.
- Goal accuracy: the agent goal accuracy metric of the RAGAS evaluation framework (version 0.4.3) (Es et al. 2024) uses an LLM judge to decide whether the conversation achieved the user’s goal (score 1) or not (score 0). As reference, the judge received the desired outcome of the question together with the result of the reference query, if available. To avoid self-evaluation, the judge model always differed from the agent model. For the comparison of the two agent models, all conversations were judged by the same model, OpenAI’s reasoning model o3 (OpenAI 2025d), which differs from both agent models. In addition, GPT-5 judged all GPT-4.1 runs and GPT-4.1 judged all GPT-5 runs, in two independent judging passes each, to assess the dependence of goal accuracy on the judge and how often two judgements of the same run agree. In the pilot run, GPT-5 agreed better with our manual assessment than GPT-4o, but it occasionally rejected correct answers.

The benchmark questions, reference queries, evaluation code, and raw results are publicly available (see Data, Code, and Application Availability).

## 4 Discussion

EcoToxFred connects complex data with practical application in environmental research by democratizing access to specialized domain knowledge. By accepting natural language queries, the system empowers environmental scientists, regulators, and the public to obtain insights without requiring computer science expertise. This lowered entry barrier fosters a more inclusive, data-driven culture in environmental decision-making, enabling diverse stakeholders to engage with evidence (e.g., chemical risk data, trends) previously inaccessible behind technical interfaces.

EcoToxFred delivers precise, context-rich answers through tabular summaries and maps, providing direct improved support for policy and risk assessment. Regulators and scientists can efficiently identify contaminated waterways with geospatial visualization of affected sites, while accessing the underlying data for verification and further analysis. This responsive analytics capability facilitates evidence-based decisions, enabling faster interventions and more informed environmental policies. In this way, raw monitoring data become actionable knowledge more quickly, which can improve both the timeliness and quality of environmental management decisions.

EcoToxFred seamlessly integrates heterogeneous knowledge by retrieving structured information from the knowledge graph (e.g., chemical occurrence counts) while incorporating contextual background from Wikipedia. This integration provides users with comprehensive answers that combine scientific context about chemicals with their environmental occurrence data. The system functions as a knowledge hub that synthesizes information across multiple sources, representing practical advancement toward AI assistants capable of reasoning across databases, documents, and ontologies simultaneously.

EcoToxFred enhances interpretability through multi-modal outputs, including interactive maps, tables, and text. By presenting results visually and in structured formats, the system helps users identify patterns, such as pollution hotspots or chemical co-occurrence relationships, that would be challenging to discern from texts alone. This approach aligns with expert analytical practices, augmenting human insight rather than simply providing textual answers. The transparency afforded by structured data presentation, such as tabular results with quantifiable metrics, enables users to verify information and builds trust in the system’s outputs. These design choices in human-AI interaction improve user comprehension of complex environmental data.

The benchmark evaluation shows to what extent EcoToxFred handles complex, multi-hop queries through query decomposition and tool selection. Lookup questions were answered reliably, whereas analytical questions that combine several conditions or aggregations succeeded in about half of the runs with both models. The errors point to concrete improvements that do not depend on the LLM. Passing more result rows, or a complete result table, to the user would close most of the gap between execution and goal accuracy. Case-insensitive matching and synonym tables for categorical properties would address the vocabulary mismatches reported in the workshop. Storing only actual driver events in the `IS DRIVER` relationship, instead of flagging non-drivers with a property, would remove a trap that misled both models. The two LLMs reached the same execution accuracy, but GPT-5 more often achieved the user’s goal, at the cost of about three times longer response times; offering both back ends allows users to choose between them. The system’s ability to generate and execute sophisticated Cypher queries, as demonstrated in the co-occurrence example, illustrates automated reasoning over the knowledge graph. This functionality positions LLM-based agents as intermediaries that translate high-level questions into database queries, which is particularly valuable in domains with complex relational data. This allows domain experts to focus on result interpretation rather than crafting technical queries. Recent research indicates that agentic LLM approaches incorporating planning steps, task decomposition, and context management significantly improve performance on complex problems (Webb et al. 2023). EcoToxFred therefore maintains an explicit to-do list for multi-step requests.

While EcoToxFred is built for ecotoxicology, the system’s generalizability allows the approach to extend to other domains. The core framework (LLM + graph + tools) could be transferred to alternative domains, which is significant for many scientific fields (medicine, biodiversity, climatology) with extensive structured databases underutilized by non-experts. EcoToxFred serves as a *blueprint* for developing similar conversational agents. For example, a “ClimateFred” could interface with climate observation databases, or a biomedical agent could query gene-disease networks. This cross-domain potential amplifies the work’s impact and aligns with a broader vision of LLMs as universal interfaces to data.

Ensuring accurate interpretation of AI outputs by users represents a significant challenge. Ecotoxicological data contains nuanced concepts (e.g., “toxic unit” definitions or detection limits) that non-experts may misinterpret without proper context. While EcoToxFred addresses this partially by providing explanations in its responses and defining terms when explicitly requested, additional user support mechanisms may be necessary. This reflects a broader challenge in scientific AI tools: balancing interface simplicity with sufficient contextual information. Potential enhancements include implementing on-demand definitions, data uncertainty indicators, and interactive tutorials to improve user comprehension and appropriate application of the information provided.

Maintaining EcoToxFred requires ongoing knowledge graph and infrastructure management to ensure reliable operation and continuous improvement. The system’s Neo4j database is hosted at the UFZ on permanently available Helmholtz infrastructure, with support from the research data management department ensuring reliable operation. The knowledge graph receives continuous updates and extensions through multiple ongoing projects. Long-term sustainability depends on implementing automated graph update mechanisms for ingesting new monitoring data and maintaining robust hosting solutions. This presents an opportunity to align with open-data initiatives within Helmholtz and the broader scientific community. Integration with regularly updated environmental databases would transform EcoToxFred into an enduring community resource with expanding utility.

The agent’s performance is directly tied to the data completeness and quality of the underlying knowledge graph. Any gaps or errors in the knowledge graph, whether missing geographical regions, species, or chemical compounds, will be reflected in the system’s responses. The current system’s focus on European surface waters represents both a limitation and an opportunity: while questions outside this scope cannot be answered, the system’s architecture allows for continuous expansion. As additional data sources covering new regions, pollutants, or biodiversity metrics are integrated, the utility and applicability of the system will proportionally increase, enhancing its overall value to the scientific community.

Handling ambiguity and query limits is addressed by the system through a feedback loop when parsing difficulties arise, either retrying or falling back to a general LLM. While this enhances robustness, certain queries remain beyond the system’s capabilities, particularly those requiring deep causal reasoning or the ones falling outside the domain scope. The system is designed to communicate uncertainty appropriately (e.g., “I’m not sure, but here is what I found …”), maintaining user trust through transparency. This approach exemplifies responsible AI design by acknowledging limitations rather than providing potentially misleading responses. Future development could incorporate user feedback mechanisms where the system requests clarification for ambiguous queries, further improving the interaction experience.

A significant technical challenge arises when database queries return large result sets, as LLMs cannot process millions of data points within their context window constraints. To address this limitation, EcoToxFred implements several complementary strategies: (1) tabular visualizations that present manageable subsets of data while providing the original Cypher query for users to conduct further exploration; (2) statistical summaries that enable the LLM to generate accurate textual descriptions from condensed data representations; and (3) specialized handling for geographical visualizations where the LLM generates the query but deterministic code manages the data processing and visualization. These approaches maintain system responsiveness while ensuring data integrity, demonstrating how hybrid architectures can overcome fundamental LLM limitations. While future models may feature expanded context windows, the principled management of large datasets will remain an essential consideration in environmental data systems.

Ensuring responsible AI and addressing potential bias are important considerations for EcoToxFred. While EcoToxFred mitigates factual accuracy issues through grounded answers, potential biases remain a consideration. Geographic biases in the underlying data may lead to systematic emphasis on certain regions, while Wikipedia snippets used for chemical information might contain phrasing that inadvertently downplays or exaggerates certain risks. The system employs extensive prompt engineering, clear agent instructions, and a zero-temperature LLM setting to ensure a balanced, scientifically accurate context. Users should nevertheless cross-verify critical information. The benchmark introduced here enables systematic regression testing when the model, prompts, or knowledge graph change; extending it with questions that probe problematic outputs, particularly regarding chemical safety information, will help to uphold *responsible AI* criteria. The current design, using curated databases and published sources, provides a foundation for responsible AI implementation, though ongoing vigilance remains essential.

The benchmark and the expert workshop provide a first quantitative and qualitative evaluation of EcoToxFred, but both have limitations. The benchmark is small, covers a single knowledge graph, and its questions and reference queries were written by the developers or derived from a single workshop, so it may not represent the full range of user questions. Goal accuracy relies on an LLM judge, which can itself err, whereas execution accuracy is strict and may count semantically equivalent answers with a different result structure as incorrect; we therefore report both. The workshop feedback was anonymous and unstructured and does not allow conclusions on time savings or on differences between user groups. Most of the obstacles reported in the workshop were not errors in query generation but mismatches between user vocabulary and the database vocabulary, such as abbreviations, chemical classes, or use classes, and concepts that are absent from the knowledge graph, such as modes of action. Adding synonym tables and mode-of-action annotations to the knowledge graph is therefore a priority. A structured user study with environmental scientists and policy analysts, including task completion times compared to conventional data analysis, is a next step. Requested features, such as data export and access to previous conversations, will guide further development.

Future development of EcoToxFred will include extended tool capabilities, such as scientific plotting capabilities and CSV export functionality, to enhance the system’s utility. Implementing a plotting tool would enable users to visualize temporal trends in chemical concentrations, enriching the interactive analysis experience. Additionally, integrating support for the recently emerged Model Context Protocol (MCP) will help EcoToxFred to produce better, more relevant responses (Anthropic 2024). These enhancements would transform EcoToxFred into a more comprehensive analysis assistant that not only answers questions but also helps visualize data patterns and comparisons dynamically. Such advancements would position the system as a collaborative research tool that augments scientific workflows rather than merely providing information.

EcoToxFred demonstrates portability and reproducibility through its publicly available codebase on GitHub. A continuous integration workflow ensures code updates are delivered to the permanent Helmholtz HIFIS GitLab service that supports the official application. The provision of a Docker image facilitates deployment and extension by other researchers on their own infrastructure. This approach aligns with modern scientific standards for transparency and reproducibility. The open architecture enables collaboration with both the AI community and domain specialists who can integrate new data or adapt the system to alternative datasets. EcoToxFred thus serves as a model for how AI-driven science communication tools can be effectively disseminated for broad scientific use.

EcoToxFred represents part of an emerging trend toward making scientific knowledge more interactive and accessible through AI. Similar to systems like ChatClimate and chemistry assistants that help navigate complex scientific information Bran et al. (2024), EcoToxFred points toward a future where environmental knowledge is readily accessible through conversational interfaces. This approach could transform science communication by enabling stakeholders to interact directly with environmental databases during public consultations, or allowing journalists to fact-check environmental claims efficiently. The potential outcome is more informed public discourse and transparent decision-making, facilitated by AI systems that make specialized data understandable to a broad audience. This work thus carries significance beyond its technical implementation, contributing to broader goals of scientific accessibility and public engagement.

## 5 Conclusion

EcoToxFred demonstrates how retrieval-augmented large language models, integrated with structured knowledge in graph databases, can significantly enhance access to complex environmental data. By connecting natural language queries with ecotoxicological data stored in a Neo4j graph database, EcoToxFred empowers researchers, policymakers, and environmental managers to engage with domain-specific knowledge without requiring expertise in database querying or data management. This lowers the barrier to working with environmental monitoring data and supports risk assessments, and therefore more informed and timely decision-making.

The knowledge graph grounding EcoToxFred originates from a separate open-science initiative and is publicly available on Zenodo, emphasizing a commitment to transparency and data reuse. Our system shows that coupling such domain-specific knowledge graphs with AI reasoning can synthesize heterogeneous data sources into coherent, actionable insights. Although challenges remain, including data completeness, user onboarding, and handling large-scale datasets, the system’s architecture is designed with scalability, transparency, and continuous improvement in mind.

Future development will focus on expanding datasets, integrating advanced analytical and visualization tools, enriching the knowledge graph with synonyms and mode-of-action annotations, and conducting structured user studies to enhance practical applicability. While rooted in ecotoxicology, this approach is transferable to other data-rich fields such as climate science, public health, and biodiversity research.

EcoToxFred thus marks a promising advance in AI-supported science communication, transforming static environmental data into dynamic, conversational insights that foster more inclusive, evidence-based environmental safekeeping.

## Data, Code, and Application Availability

The underlying Neo4j graph database container that stores the structured ecotoxicological data is published separately and described in Schor et al. (2025a). The source code for EcoToxFred, including the conversational agent implementation, data retrieval modules, and interaction interfaces, is publicly available on GitHub: https://github.com/yigbt/EcoToxFred. The fully functional web application is accessible at https://ecotoxfred.web.app.ufz.de/. The benchmark questions, reference queries, evaluation code, and raw benchmark results are part of the repository (directory `benchmark`) and archived at Zenodo with release 2026.1.0 (Scheibe et al. 2026). All resources are openly licensed and designed for reuse and extension by the scientific community.

## Supplementary information

Supplementary Table S1 lists all benchmark questions with their category, origin, expected tools, reference query, and per-question results.

## CRediT authorship contribution statement

Patrick Scheibe: conceptualization, methodology, software, writing – original draft, writing – review & editing. Thilo Simon Schmid: software, methodology, validation, writing – review & editing. Jana Schor: conceptualization, methodology, software, data curation, validation, formal analysis, investigation, writing – original draft, writing – review & editing, supervision, project administration, funding acquisition.

## Declaration of competing interest

The authors declare that they have no known competing financial interests or personal relationships that could have appeared to influence the work reported in this paper.

## Declaration of generative AI and AI-assisted technologies in the writing process

During the preparation of this work, the authors used Claude (Anthropic) to assist with revising the manuscript text and implementing the benchmark evaluation code. After using this tool, the authors reviewed and edited the content as needed and take full responsibility for the content of the publication. EcoToxFred itself uses LLMs as described in the Methods section.

## Acknowledgments

We thank the participants of the workshop “CITE Meets AI: Working with LLMs, EcoToxFred, and WISE” for testing EcoToxFred and for their feedback. We gratefully acknowledge the UFZ Research Data Management (RDM) unit for their support in deploying the Neo4j container, setting up the EcoToxFred web application, and ensuring seamless integration between the two components.

## Funding

We acknowledge financial support from the Helmholtz POF IV Topic 9, Germany: “Healthy Planet– towards a non-toxic environment.” This work was also supported by the Partnership for the Assessment of Risks from Chemicals (PARC), a European research project involving nearly 200 institutions from 28 countries and three EU authorities on chemical risk assessment; The European Partnership for the Assessment of Risks from Chemicals has received funding from the European Union’s Horizon Europe research and innovation programme under Grant Agreement No. 101057014. Views and opinions expressed are, however, those of the author(s) only and do not necessarily reflect those of the European Union or the Health and Digital Executive Agency. Neither the European Union nor the granting authority can be held responsible for them.

## Notes

https://ecotoxfred.web.app.ufz.de/

https://github.com/yigbt/EcoToxFred

### Competing Interest Statement

The authors have declared no competing interest.

### Summary of Updates

This version adds a quantitative evaluation of EcoToxFred: a benchmark of 25 questions with human-written reference queries, evaluated for tool selection, execution accuracy, and goal accuracy (RAGAS) with GPT-4.1 and GPT-5 as agent models, and an analysis of anonymous feedback from an expert workshop, six failure cases of which were turned into benchmark questions. The Methods describe the current release of EcoToxFred (2026.1.0, archived at Zenodo, https://doi.org/10.5281/zenodo.24219536). The title and abstract were revised, Thilo Simon Schmid joined as an author, and Supplementary Table S1 lists all benchmark questions and per-question results.

