## Supplemental material for "EcoToxFred: A knowledge-graph-grounded conversational agent for exploring ecotoxicological monitoring and hazard data"

### Supplementary Table S1: Benchmark questions and per-question results

Each question was posed three times per agent model. T: tool selection; EX: execution accuracy; G: goal accuracy (RAGAS, judged by o3 for both models); all values in percent of runs. EX is not assessed (–) for open-ended questions. D: questions written by the developers; W: questions derived from the workshop feedback. Reference Cypher queries, answer columns, desired outcomes, full conversations, and raw scores are available in the `benchmark` directory of the EcoToxFred repository.

| | | | | GPT-4.1 | | | GPT-5 | | |
|---|---|---|---|---|---|---|---|---|---|
| ID | Category | Question | Expected tools | T | EX | G | T | EX | G |
| D00 | General | What is DDT and why is it harmful to the environment? | WikipediaSearch | 100 | – | 100 | 100 | – | 100 |
| D01 | Analytical | Find the 10 most frequent multiple risk drivers | CypherSearch | 100 | 0 | 0 | 0 | 0 | 0 |
| D02 | Lookup | Which countries have the highest number of sampling sites in the dataset, and how many sites does each of the top 10 countries contain? | CypherSearch | 100 | 100 | 33 | 100 | 100 | 67 |
| D03 | Lookup | What is the temporal coverage (min/max year) of the monitoring data stored on MEASURED_AT relationships? | CypherSearch | 100 | 100 | 100 | 100 | 100 | 100 |
| D04 | Lookup | What is the temporal coverage (min/max year) of mixture toxicity summaries stored on SUMMARIZED_IMPACT_ON relationships? | CypherSearch | 100 | 100 | 100 | 100 | 100 | 100 |
| D05 | Lookup | For Diuron, at how many sites was it detected (defined as median_concentration > 0) across all years/quarters? | CypherSearch | 100 | 100 | 100 | 100 | 100 | 100 |
| D06 | Lookup | Which sites have the highest number of distinct detected substances (unique Substance nodes with median_concentration > 0) aggregated over all sampled year-quarters? | CypherSearch | 100 | 100 | 0 | 100 | 100 | 0 |
| D07 | Analytical | For each species group (algae, crustacean, fish), which countries have the highest number of site-year-quarter events with a sumTU > 1 (acute mixture toxicity)? | CypherSearch | 100 | 100 | 67 | 100 | 0 | 33 |
| D08 | Analytical | For each species group (algae, crustacean, fish), which country has the highest number of site-year-quarter events with a sumTU > 0.001 (chronic mixture toxicity)? | CypherSearch | 100 | 100 | 100 | 100 | 100 | 100 |
| D09 | Analytical | At site-year-quarter resolution, what fraction of driver events (IS_DRIVER with is_driver = true) coincide with a detection of the same substance (median_concentration > 0 on MEASURED_AT) at the same site, year and quarter, and does this fraction differ by species? | CypherSearch | 100 | 100 | 100 | 100 | 100 | 100 |

| | | | | GPT-4.1 | | | GPT-5 | | |
|---|---|---|---|---|---|---|---|---|---|
| ID | Category | Question | Expected tools | T | EX | G | T | EX | G |
| D10 | Analytical | For each species, which substance reaches TU > 1 most often (acute toxicity at the single-substance TU level via TU_algae/TU_crustacean/TU_fish on MEASURED_AT), and at how many distinct sites do these exceedances occur? | CypherSearch | 100 | 0 | 0 | 100 | 33 | 100 |
| D11 | Analytical | For each species, which substances most often exceeds TU > 0.001 (potential chronic impacts), which substances are in the top 1% for all three species? | CypherSearch | 100 | – | 33 | 67 | – | 33 |
| D12 | Analytical | For Algae: at events with sumTU > 1, how many substances typically contribute TU > 0.001 at that same site-year-quarter? | CypherSearch | 100 | 67 | 0 | 100 | 33 | 67 |
| D13 | Analytical | For each species return the avg ratioTU and the 25th and 75th percentile of ratios across all sites? | CypherSearch | 100 | 33 | 0 | 100 | 100 | 0 |
| D14 | Analytical | For site-year-quarter events where the sumTU for both algae and fish exceeds 0.0001, what is the average ratio of the fish sumTU to the algae sumTU, and what are the 2.5th and 97.5th percentiles of this ratio? | CypherSearch | 100 | 33 | 33 | 100 | 67 | 100 |
| D15 | Analytical | Which river basins (Site.river_basin) show the highest rates of (a) detected substances, (b) driver substances, and (c) per species sumTU exceedances, and how do these metrics compare across basins? | CypherSearch | 100 | – | 0 | 67 | – | 0 |
| D16 | Analytical | Which pairs of substances have the highest co-driver frequency (via JOINT_DRIVER_WITH relationship frequency), and do those substances also tend to be detected at the same sites? | CypherSearch | 100 | – | 100 | 100 | – | 100 |
| D17 | Analytical | Are there countries/river basins where the driver network (substances linked by JOINT_DRIVER_WITH) is denser (more high-frequency co-driver edges), compared to others? | CypherSearch | 100 | – | 100 | 67 | – | 100 |
| D18 | Analytical | Using TESTED_FOR_TOXICITY relationships (tox_value_mg_L) and measured concentrations (MEASURED_AT), which substances have the smallest margin between measured levels and toxicity values (i.e., highest implied risk), and does this align with them being labeled drivers (IS_DRIVER)? | CypherSearch | 0 | – | 0 | 100 | – | 67 |
| W01 | Workshop | In which sites was Iomeprol found to be a risk driver in Germany in 2010? | CypherSearch, GeographicMap | 100 | 100 | 33 | 100 | 100 | 33 |
| W02 | Workshop | In which sites was Iomeprol found to be a risk driver in France in 2010? | CypherSearch, GeographicMap | 100 | 100 | 100 | 100 | 67 | 100 |
| W03 | Workshop | What are the most frequent risk drivers for species algae that fall into the use class of pharmaceuticals? | CypherSearch | 100 | 0 | 0 | 100 | 33 | 0 |
| W04 | Workshop | List me 20 chemicals which are more than 10x more toxic in fish than in algae and daphnia. | CypherSearch | 100 | 100 | 100 | 100 | 100 | 67 |

| | | | | GPT-4.1 | | | GPT-5 | | |
|---|---|---|---|---|---|---|---|---|---|
| ID | Category | Question | Expected tools | T | EX | G | T | EX | G |
| W05 | Workshop | Which estrogenic compounds have been measured, and where? | CypherSearch, WikipediaSearch | 100 | – | 0 | 100 | – | 0 |
| W06 | Workshop | At how many sites has DDT been measured? | CypherSearch | 100 | 100 | 100 | 100 | 100 | 100 |
